# Ethanolamine-induced assembly of microcompartments is required for *Fusobacterium nucleatum* virulence

**DOI:** 10.1101/2024.11.11.623001

**Authors:** Dana S. Franklin, Yi-Wei Chen, Yimin Chen, Manuel Wittchen, Angela Agnew, Alexis Luu, Julian P Whitelegge, Z Hong Zhou, Andreas Tauch, Asis Das, Hung Ton-That

## Abstract

Many bacteria metabolize ethanolamine as a nutrient source through cytoplasmic organelles named bacterial microcompartments (BMCs). Here we investigated the molecular assembly, regulation, and function of BMCs in *Fusobacterium nucleatum* – a Gram-negative oral pathobiont that is associated with adverse pregnancy outcomes. The *F. nucleatum* genome harbors a conserved ethanolamine utilization (*eut*) locus with 21 genes that encode several putative BMC shell proteins and a two-component signal transduction system (TCS), in addition to the enzymes for ethanolamine transport and catabolism. We show that the expression of most of these genes as well as BMC formation is highly increased in wild type fusobacteria when cultured in the presence of ethanolamine as a nutrient source. Deletion of the response regulator EutV eliminated this induction of *eut* mRNAs and BMCs, thus demonstrating that BMC formation is transcriptionally regulated by the TCS EutV-EutW in response to ethanolamine. Mass spectrometry of isolated BMCs unveiled the identity of the constituent proteins EutL, EutM_1_, EutM_2_, and EutN. Consistent with the role of these proteins in BMC assembly and metabolism, deletion of *eutN*, *eutL*/*eutM_1_*/*eutM_2_*, or *eutL*/*eutM_1_*/*eutM_2_*/*eutN* not only affected BMC formation, but also ethanolamine utilization, causing cell growth defects with ethanolamine as nutrient. BMCs also assembled in fusobacteria cultured with placental cells or the culture media, a process that is dependent on the BMC shell proteins. Significantly, we show that the *eutN* mutant is defective in inducing preterm birth in a mouse model. Together, these results establish that BMC-mediated metabolism of ethanolamine is critical for fusobacterial virulence.

**IMPORTANCE:** The oral anaerobe *Fusobacterium nucleatum* can spread to distal internal organs, such as the colon and placenta, and thereby promote the development of colorectal cancer and induce preterm birth, respectively. Yet, how this opportunistic pathogen adapts to the various metabolically distinct host cellular niches remains poorly understood. We demonstrated here that this microbe assembles specialized metabolic organelles, termed bacterial microcompartments (BMCs), to utilize environmental ethanolamine (EA) as a key environmental nutrient source. The formation of *F. nucleatum* BMCs, containing BMC shell proteins EutLM_1_M_2_N, is controlled by a two-component system, EutV-EutW, responsive to EA. Significantly, this ability of *F. nucleatum* to form BMCs in response to EA is crucial for its pathogenicity evidenced by the fact that the genetic disruption of BMC formation reduces fusobacterial virulence in a mouse model of preterm birth.

## INTRODUCTION

Ethanolamine (EA) – a component of the membrane lipid phosphatidylethanolamine present in both eukaryotes and prokaryotes – is a notable nutrient source to gut bacteria, such as *Escherichia*, *Salmonella*, and *Listeria* (1, 2). Each of these microbes harbor a multi-gene locus, named *eut* for <u>e</u>thanolamine <u>ut</u>ilization, that encodes various proteins for EA transport and metabolism (1, 2). The *eut* locus also encodes several small structural proteins known as shell proteins (3). These shell proteins self-assemble to form polyhedral organelles called the bacterial microcompartments (BMCs) that encapsulate the EA catabolic enzymes and thereby empower bacteria with a highly efficient catabolic machine while also compartmentalizing and insulating bacterial cells from the toxic intermediates of EA catabolism (4, 5). In *Escherichia coli*, BMCs were shown to contain EutS, EutL, EutK, EutM, and EutN, with the first four representing the major shell constituents (6). Importantly, genetic studies in *Salmonella enterica* revealed that individual deletion mutants of *eutK*, *eutL*, *eutM*, or *eutN* fail to grow on EA as a carbon source at high pH (≥ 8), whereas mutant cells lacking *eutS* grow similarly as wild-type cells (7). Expression of the *S. enterica* shell proteins EutSMNLK in the heterologous host *E. coli* is sufficient to produce *Salmonella* BMCs, with increased BMC formation when EutQ is co-expressed (8). However, whether mutants lacking the aforementioned individual shell proteins are defective in BMC formation remained unclear.

Two distinct types of regulatory systems are known to be involved in controlling the formation of BMCs and the expression of *eut* genes in different organisms – the transcriptional regulator EutR and the two-component transduction system (TCS) EutV-EutW, with EutV acting as a response regulator and EutW as a sensor histidine kinase (1). BMCs are observed in wild-type *Listeria monocytogenes* grown in the EA-containing media but not in mutants lacking *eutV* (9). In contrast, EutR was shown to be a direct regulator of the *eut* locus in enterohemorrhagic (i) *E. coli* (EHEC), as EutR binds to the promoter of *eutS* – the first gene of the *eut* locus, and *eutR* deletion drastically reduces *eutS* expression (10).

The multigene *eut* locus is present in the genomes of a wide range of Gram-positive and Gram-negative bacteria, including the subject of this study, *Fusobacterium nucleatum*, whose genome harbors three short operons of roughly 17 *eut* genes (3, 11, 12). *F. nucleatum*, a Gram-negative obligate anaerobe, is mainly found in the human oral cavity, but it has the ability to spread to distal organs such as colon and placenta (13–19), where the bacterium was found to promote colorectal cancer and adverse pregnancy outcomes including neonatal sepsis and preterm birth, respectively (20). The *eut* gene cluster is conserved in many other *F. nucleatum* subspecies such as *F. nucleatum* ssp. *vincentii* and *F. nucleatum* ssp. *animalis* (12, 21, 22). Consistent with the functionality of the *eut* gene locus in *F. nucleatum*, *F. nucleatum* ssp. *animalis* cells exposed to EA display a significant increase in expression of *eut* genes including those that encodes BMC proteins (22). This conservation of the EA utilization system and its regulation by EA and BMCs in a pathologically important fusobacterial clade underscores the importance of dissecting the genetic, molecular, and physiological determinants of BMC assembly and the role of BMCs in fusobacterial pathogenicity.

Here, using the *F. nucleatum* sub spp. *nucleatum* strain ATCC 23726, a clinical isolate that is amendable to robust genetic manipulation (23–28), we show that fusobacteria are responsive to EA as an environmental nutrient source that greatly increases the expression of *eut* genes and BMC formation. This EA response of fusobacteria requires the response regulator EutV, which appears to be an activator of *eut* genes and also a global transcriptional regulator of a large regulon under nutrient-rich conditions. We further show that *F. nucleatum* BMCs contain EutL, EutM_1_, EutM_2_, and EutN, and genetic disruption of *eutN*, *eutLM_1_M_2_*, or *eutLM_1_M_2_N* severely affects BMC formation in EA-supplementing conditions as well as in co-cultures with placental cells. Most importantly, we show that the BMCs are essential for fusobacterial pathogenesis by demonstrating that the *eutN* mutant is significantly attenuated in virulence in a mouse model of preterm birth.

## RESULTS

### The conserved *eut* locus in *F*. *nucleatum* responds to environmental ethanolamine

To elucidate the functionality of the *eut* gene cluster in *F. nucleatum*, we chose a genetically tractable clinical isolate of *F. nucleatum* ssp. *nucleatum*, ATCC 23726. The genome of this strain contains three short operons within a conserved gene cluster with several sets of genes that are characteristic of the *eut* locus: a two-component transduction system (TCS) (EutV-EutW), several putative bacterial microcompartment (BMC) proteins (EutS-EutL-EutM), an EA transporter (EutH), EA catabolic proteins (EutA/EutB/EutC) (Fig. 1A). The cluster also encodes several hypothetical proteins (RS02810 and RS02820), among many other predicted ethanolamine utilization proteins (EutP, EutE, EutT, EutQ, EutG, and HAD – haloacid dehalogenase-like hydrolase) (Fig. 1A). Another predicted gene, *eutJ*, coding for an EA utilization protein, is located elsewhere in the chromosome. Intriguingly, ATCC 23726 harbors 2 copies of EutM (M1 and M2) within the *eut* gene cluster, and it does not appear to encode a EutK homolog, (Fig. 1A & Fig. S1). Remarkably, this same gene arrangement was found in *F. nucleatum* ATCC 25586, as well as in many other *F. nucleatum* subspecies, including *F. nucleatum* ssp. *polymorphum* (*Fnp* F0401), *F. nucleatum* ssp. *vincentii* (*Fnv* 3_1_27), and *F. nucleatum* ssp. *animalis* (*Fna* 11_3_2) (Fig. 1A) (12), suggesting that EA utilization is a conserved feature in *F. nucleatum*.

**Figure 1:**
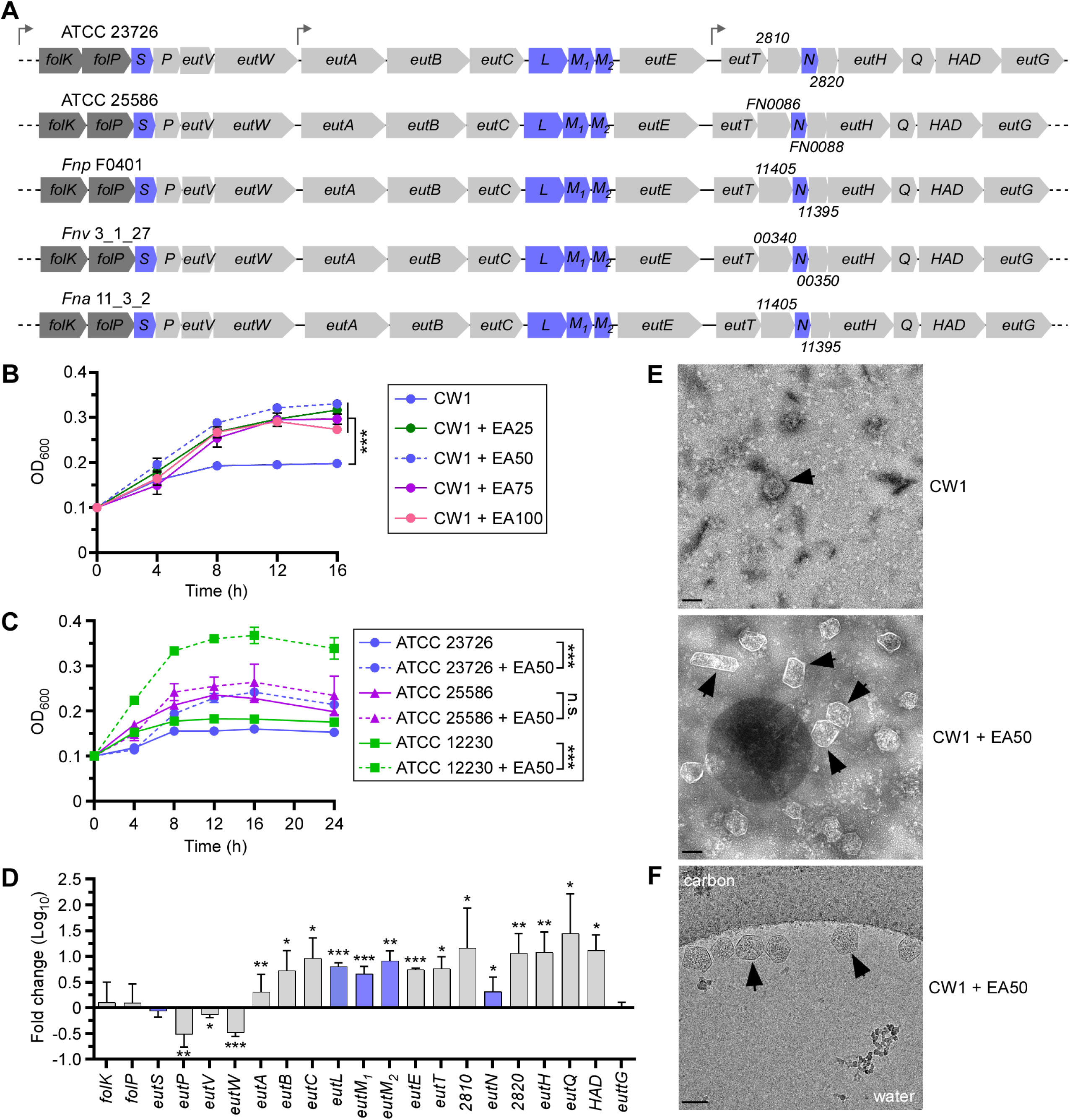
The *eut* locus of *F. nucleatum* and cellular response to ethanolamine. **(A)** Shown are the *eut* loci in *F. nucleatum* strains ATCC 23726 and 25586, with genes coding for presumptive BMC shell proteins highlighted in blue. **(B)** Cells of CW1 (derivative of ATCC 23726) were grown at 37°C in minimal media supplemented with various concentrations of EA (0, 25, 50, 75, 100 mM). Cell growth was monitored by optical density at 600 nm (OD_600_). Error bars indicate means and standard deviations of three biological replicates; ***, P<0.001. **(C)** Cells of indicated strains were grown at 37°C in minimal media supplemented with 50 mM of EA, and growth was monitored by OD_600_. Error bars indicate means and standard deviations of three biological replicates; ***, P<0.001; n.s., not significant. **(D)** CW1 cells were grown at 37°C in minimal media supplemented with 50 mM of EA for 8 h and harvested for total RNA extraction. Expression of indicated genes was determined by qRT-PCR, with 16S RNA as control. Error bars indicate means and standard deviations of three biological replicates; *, P<0.05; **, P<0.01; ***, P<0.001. **(E)** BMCs of the parent strain (CW1) grown in conditions described in B in the absence or present of 50 mM EA were isolated and analyzed by electron microscopy; scale bars of 100 nm. **(F)** Aliquots of BMCs isolated in E were analyzed by cryo-electron microscopy at the nominal magnification of 22,000 x, showing BMCs clustered along the edge of a vitreous ice hole within the holey carbon grid; scale bars of 100 nm. Black arrows in panels E and F mark representative BMC structures.

To examine how *F. nucleatum* responds to EA, we grew cultures of *F. nucleatum* CW1, an isogenic Δ*galK* derivative of ATCC 23726 that has been used for facile gene deletion (27), in minimal medium supplemented with increasing concentrations of EA, and monitored cell growth over time by measuring optical density (OD of 600 nm). *F. nucleatum* cells proliferated better and faster in the presence of EA, optimally at 50 mM (Fig. 1B; dashed blue line), compared to a culture without EA, whose growth arrested around 8 h in culture (Fig. 1B; solid blue line). To confirm that this growth phenotype is not strain-specific, we monitored the cell growth of strains *F. nucleatum* ATCC 23726, *F. nucleatum* ATCC 25586, and *F. nucleatum* ATCC 12230 in minimal medium with or without 50 mM EA. All three strains displayed a similar trend of increasing cell growth in the presence of EA, albeit no significant difference observed with ATCC 25586, with ATCC 12230 showing the largest increase (Fig. 1C). Next, we isolated total mRNA of CW1 grown in the same media in the presence or absence of 50 mM EA for 8 h for gene expression analysis by quantitative reverse transcription polymerase chain reaction (qRT-PCR). Strikingly, while the expression of a few genes of the locus remained unchanged (*eutG*, *eutS*, *eutV*), and the expression of a few others decreased (*eutP* and *eutW*), a significant increase in expression was observed for a large number of genes including those predicted to encode the BMC shell proteins as well as EA transport and catabolic enzymes (Fig. 1D). Noticeably, this bimodal expression of *eut* genes in strain 23726 was also observed in strain 25586 (Fig. S2). It is noteworthy that a similar expression pattern for many *eut* genes in *F. nucleatum* ssp. *animalis* in response to EA was also observed (22).

To investigate whether the increased expression of predicted BMC proteins caused by EA led to increased BMC formation, we isolated BMCs from fusobacterial cells grown in two conditions as performed in the previous experiment (Fig. 1D) and quantified BMCs by electron microscopy (see Methods). Indeed, as shown in Fig. 1E, transmission electron microcopy (TEM) revealed a substantial increase in the number of polyhedral BMCs in the presence of EA. These polyhedral structures were also confirmed by cryogenic-electron microscopy (cryoEM) (Fig. 1F). Together, the results suggest that *F. nucleatum* harbors a functional ethanolamine utilization system involving BMCs that is responsive to environmental EA.

### Modulation of *eut* gene expression by the response regulator EutV

The bimodal expression of *eut* genes in the presence of EA, as demonstrated above, suggests that expression of *eut* genes is regulated, most likely by the TCS EutV-EutW given their close proximity with most other *eut* genes (Fig. 1A). To examine if this is the case, we generated an in-frame, non-polar deletion mutant devoid of the predicted response regulator EutV. Next, we isolated total mRNA of the parent and *eutV* mutant strains grown in minimal medium in the presence of 50 mM EA for 8 h and quantified gene expression by qRT-PCR. Significantly, mRNAs for all the genes whose expression was greatly elevated by EA were down-regulated in the *eutV* deletion mutant, while mRNAs for a few other genes (*eutS*, *eutW*, and *eutJ*) were up-regulated when *eutV* was absent (Fig. 2A), consistent with the bimodal expression of *eut* genes as observed above (Fig. 1D). Together these results support the view that the TCS gene products EutV-EutW responds to EA, and that EutV acts as an activator of a set of *eut* genes in response to EA.

**Figure 2:**
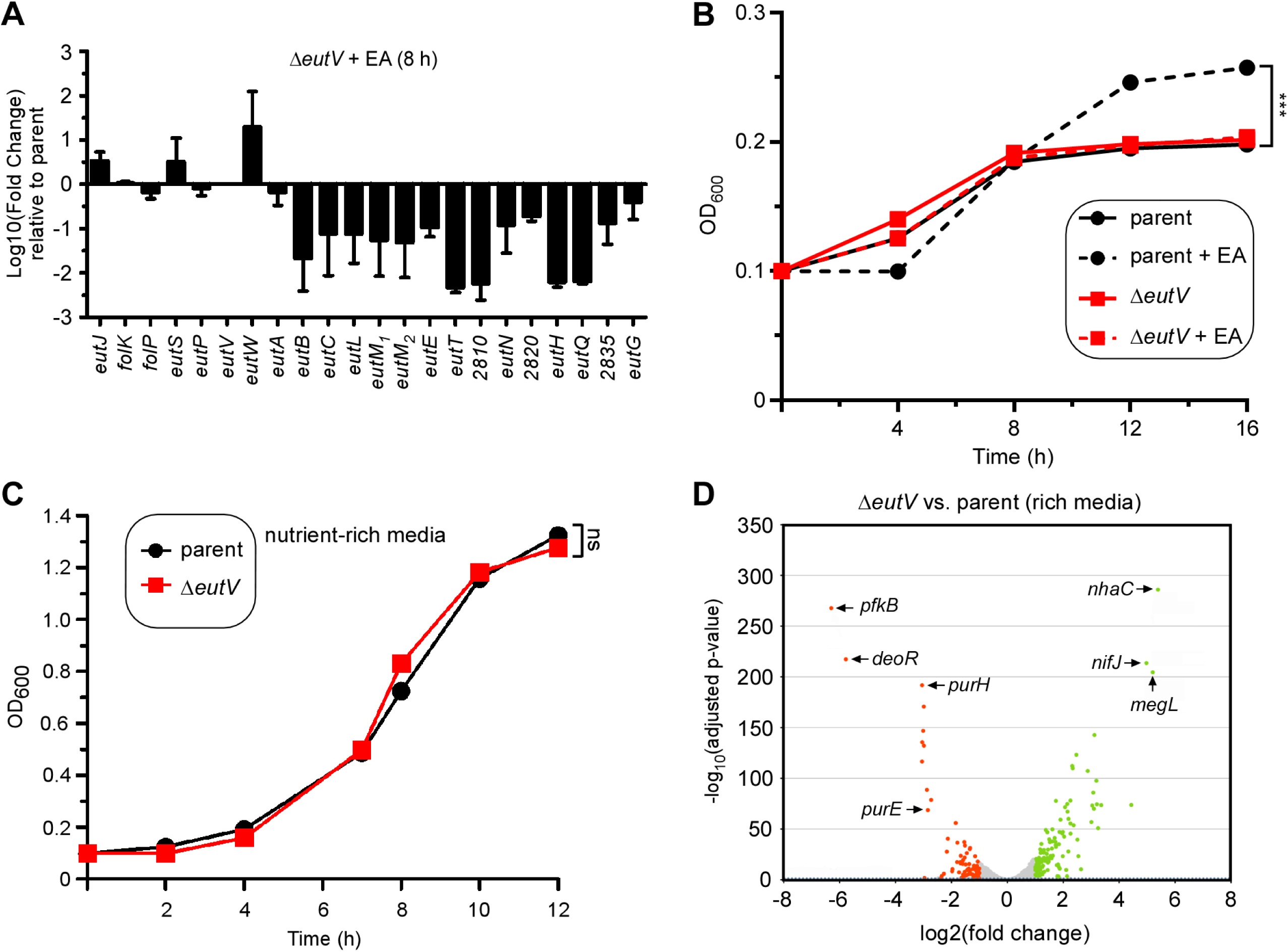
Gene expression modulated by the response regulator EutV. **(A)** Total RNA of the parent and *eutV* mutant strains grown at 37°C in minimal media supplemented with 50 mM of EA for 8 h was isolated for gene expression analysis by qRT-PCR. Error bars indicate means and standard deviations of three biological replicates. **(B)** Growth of indicated strains in minimal media supplemented with 50 mM of EA or without at 37°C was monitored by OD_600_. Error bars indicate means and standard deviations of three biological replicates; ***, P<0.001. **(C)** Total RNA of the parent and *eutV* mutant strains grown in rich media at 37°C to mid-log phase was isolated for RNA-seq analysis as previously described (25). The log_2_ (fold change) (LFC) was plotted against the statistical significance (-log_10_ of the adjusted p-value). All genes above a LFC of +1.0 and below 1.0 are shown in green and red, respectively. An insert shows a detailed portion of the volcano plot for a better presentation of the position of the adjusted p-value of 0.05 (dashed blue line). Some of the highly expressed genes are marked.

Given that many EA metabolic and BMC-encoding genes are down-regulated in the absence of EutV, the *eutV* mutant might exhibit a defect in EA utilization and cell growth when cells are grown in media with EA as the major carbon and nitrogen source. Indeed, compared to the parent strain CW1, which was positively responsive to EA for rapid growth, the *eutV* mutant displayed significant growth reduction despite the presence of EA (Fig. 2B). In contrast, both parent and Δ*eutV* mutant cells grew at similar rate in nutrient-rich media (Fig. 2C), further supporting that the loss of this regulator impacts cellular response to environmental EA.

Considering EutV regulates the *eutJ* gene that is not located at the large *eut* gene cluster, we examine whether EutV regulates additional genes of the fusobacterial genome, we performed a comparative transcriptome analysis using RNA-sequencing (RNA-seq) with total RNA isolated from the parent and *eutV* mutant strains grown to mid-log phase in rich media as previously reported (25). When a twofold cutoff (log2 fold change ± 1) was set for comparison of RNA-seq reads, we found that mRNA counts for 121 genes were significantly elevated, while that for 83 genes were significantly reduced in the absence of *eutV* (Fig. 2C and Table S1). It is not surprising that *eut* genes are not found in the list of 204 differentially expressed genes as the culture condition used for this analysis did not require EA utilization (Table S1). Noticeably, many genes up-regulated in the absence of *eutV* encode proteins predicted to be involved in sugar/amino acid metabolism and transport (Table S1), with *megL* (> 36-fold increase) coding for the methionine gamma-lyase MegL that was previously shown to participate in cysteine metabolism in *F. nucleatum* (23). On the other hand, genes coding for glycolysis (PfkB), purine biosynthesis (PurE, PurH), and methionine metabolism/transport are among the highly down-regulated genes in the absence of *eutV*. Interestingly, many genes coding for multidrug transporters (RS07785, RS07780, RS01595, RS01290, and RS10825) are also significantly down-regulated in the *eutV* mutant (Table S1). Together, the transcriptome data combined with the quantitative RNA analysis demonstrates that EutV is a response regulator that not only modulates *F. nucleatum eut* genes in response to environmental EA but also has a broad physiological role under nutrient-rich conditions in regulating the expression of many metabolic genes outside of the *eut* gene locus.

### Genetic determinants of BMC formation in *F. nucleatum*

As we alluded in the introduction, the BMCs of *Escherichia coli* and *Salmonella enterica* are made of EutS, EutK, EutL, EutM, and EutN (1). By comparison, the *eut* gene loci in *F. nucleatum* strains ATCC 23726 and ATCC 25586 do not appear to encode EutK, but instead encode two copies of EutM, i.e. EutM_1_ and EutM_2_ (Fig. 1A). Because of amenability of in ATCC 23726 for genetic manipulation (25–27), we focused on this strain to examine whether these predicted genes participate in BMC formation by generating in-frame, non-polar deletion mutants according to our published protocol (26, 29). In this method, homologous exchange of the targeted genes with deletion alleles via sequential recombination at the flanking homology regions readily produce pairs of strains with either wild-type or mutant alleles (see Methods). While we succeeded generating individual mutants devoid of *eutN*, *eutL*/*eutM_1_*/*eutM_2_*, or *eutL*/*eutM_1_*/*eutM_2_*/*eutN*, we could not obtain any mutant alleles of *eutS* after several attempts, suggesting that the deletion of this gene might be lethal in *F. nucleatum* for unknown reasons.

Next, to visualize BMCs by TEM, the parent and generated mutant strains were cultured in minimal media supplemented with 50 mM EA or without, and the harvested cells were used to isolate BMCs, which were analyzed by electron microscopy (see Methods). Compared to the parent strain grown in the presence of EA, deletion of *eutN*, *eutL*/*eutM_1_*/*eutM_2_*, or *eutL*/*eutM_1_*/*eutM_2_*/*eutN* significantly reduced BMC formation, and the resulting BMCs were visibly smaller in size (Fig. 3A). Importantly, the ectopic expression of *eutN* rescued the defect of the Δ*eutN* mutant (Fig. 3A and Fig. S3A). Furthermore, consistent with the role of BMCs in EA metabolism, all three shell mutant strains failed to respond to EA, unlike the parent strain (Fig. 3B). Note that complementation of the Δ*eutN* mutant slowly restored cell growth, which was more pronounced after 12 h (Fig. 3C).

**Figure 3:**
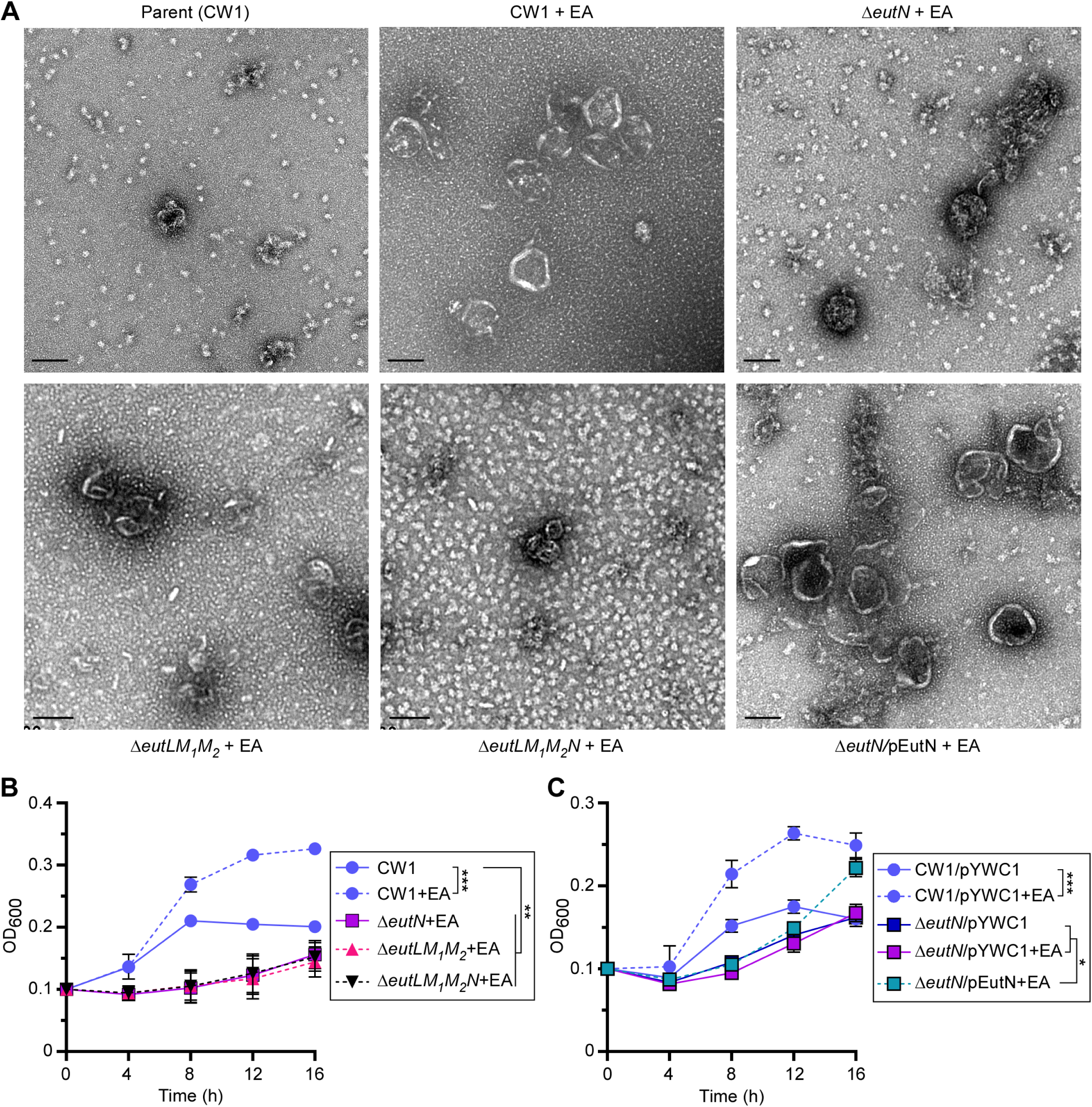
Genetic determinants of BMC formation required for selective growth. **(A)** BMCs of cells of the parent and indicated mutant strains grown in conditions described in 1E were isolated and analyzed by electron microscopy; scale bars of 100 nm. **(B-C)** Cell growth of indicated strains were monitored by OD_600_. Error bars indicate means and standard deviations of three biological replicates; *, P<0.05; **, P<0.01; ***, P<0.001.

To further confirm that EutL, EutM_1_, EutM_2_, and EutN are part of the *F. nucleatum* BMCs, we isolated BMCs from the parent strain grown in minimal media supplemented with EA for 8 h and performed mass spectrometry analysis for protein identification. As expected, we identified all aforementioned proteins among many other EA metabolic proteins, but surprisingly we did not find any signal for EutS (Table S2).

We next examined BMCs within fusobacterial cells by thin-section TEM, whereby *F. nucleatum* cells grown in the same conditions as described in Fig. 3A were harvested for preparation of thin-sections, which were stained with 1% uranyl acetate prior to TEM observation (see Methods). Consistent with the above results, the parent cells (CW1) produced a higher number of BMCs in the presence of 50 mM EA than without it (Fig. 4A-4B). Significantly, all three shell protein mutants (single, triple, and quadruple) produced less BMCs than the parent strain (Fig. 4C-4E), whereas complementation of the Δ*eutN* mutant rescued this defect (Fig. 4F and Fig. S3B).

**Figure 4:**
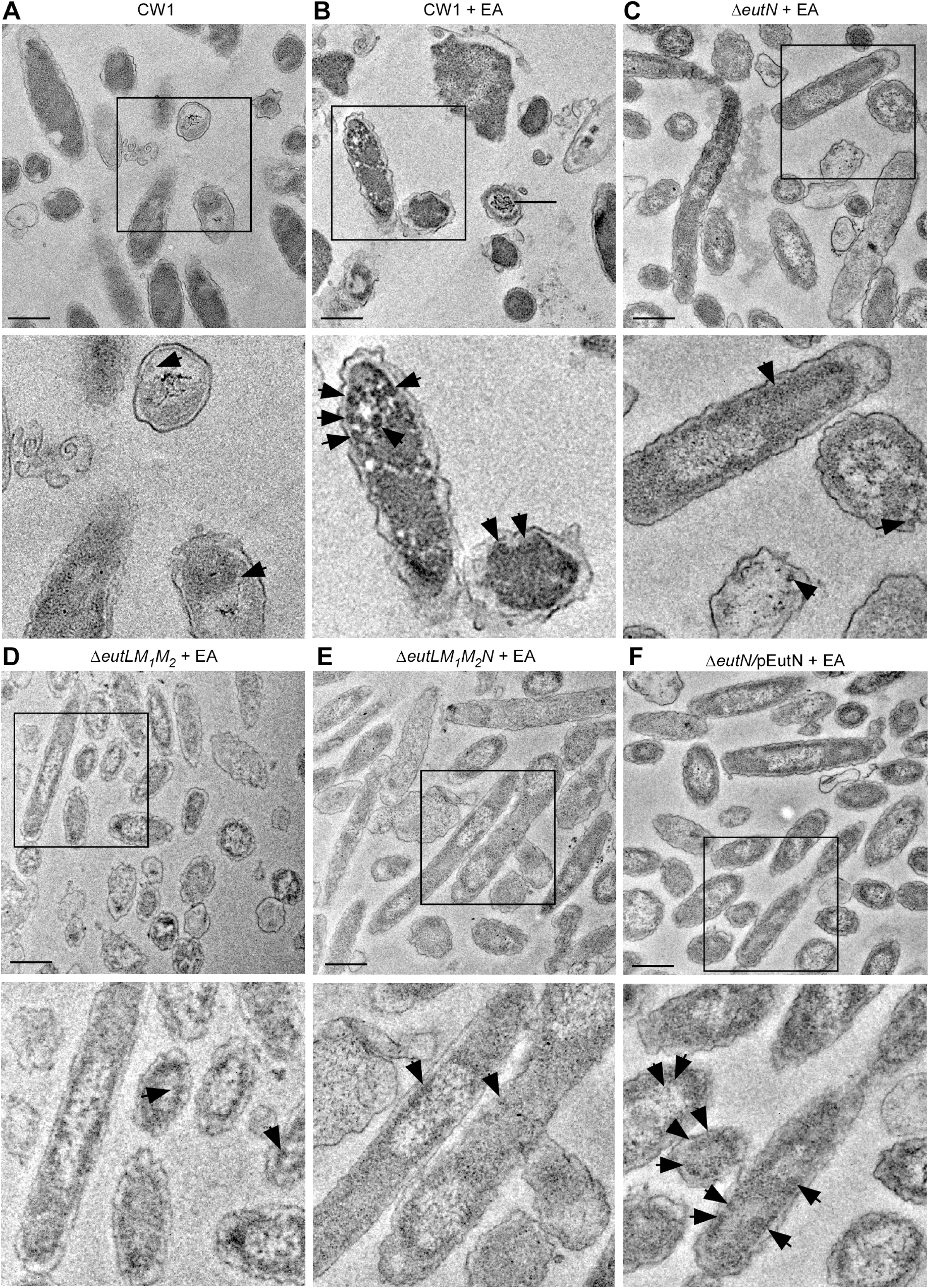
BMC formation analyzed by thin-section electron microscopy. **(A-F)** Cells of the parent and indicated mutant strains grown in conditions described in 1E were harvested for thin-section electron microscopy; scale bars of 200 nm. Enlargement of boxed areas is shown in below panels. Black arrows mark representative BMC structures.

Finally, we examined whether *F. nucleatum* forms BMCs in culture in the presence of placental cells. We co-cultured fusobacterial cells with HTR8/SVneo placental cells for 4.5 h prior to embedding them in resin for thin-sectioning. Compared to the parent strain, which formed BMCs in the presence of HTR8, the Δ*eutN* mutant produced significantly less BMCs (Fig. 5A-5B). This defect was recused by ectopic expression of *eutN* (Fig. 5C). Notably, BMCs were rarely observed in the triple and quadruple mutants (Fig. 5D-5E). Interestingly, the parent CW1 cells grown in media used for growing HTR8 cells (RPMI+ 5% FBS) produced a high number of BMCs (Fig. 5F and Fig. S3C), suggesting these media contain EA and/or some other inducer that triggered BMC formation.

**Figure 5:**
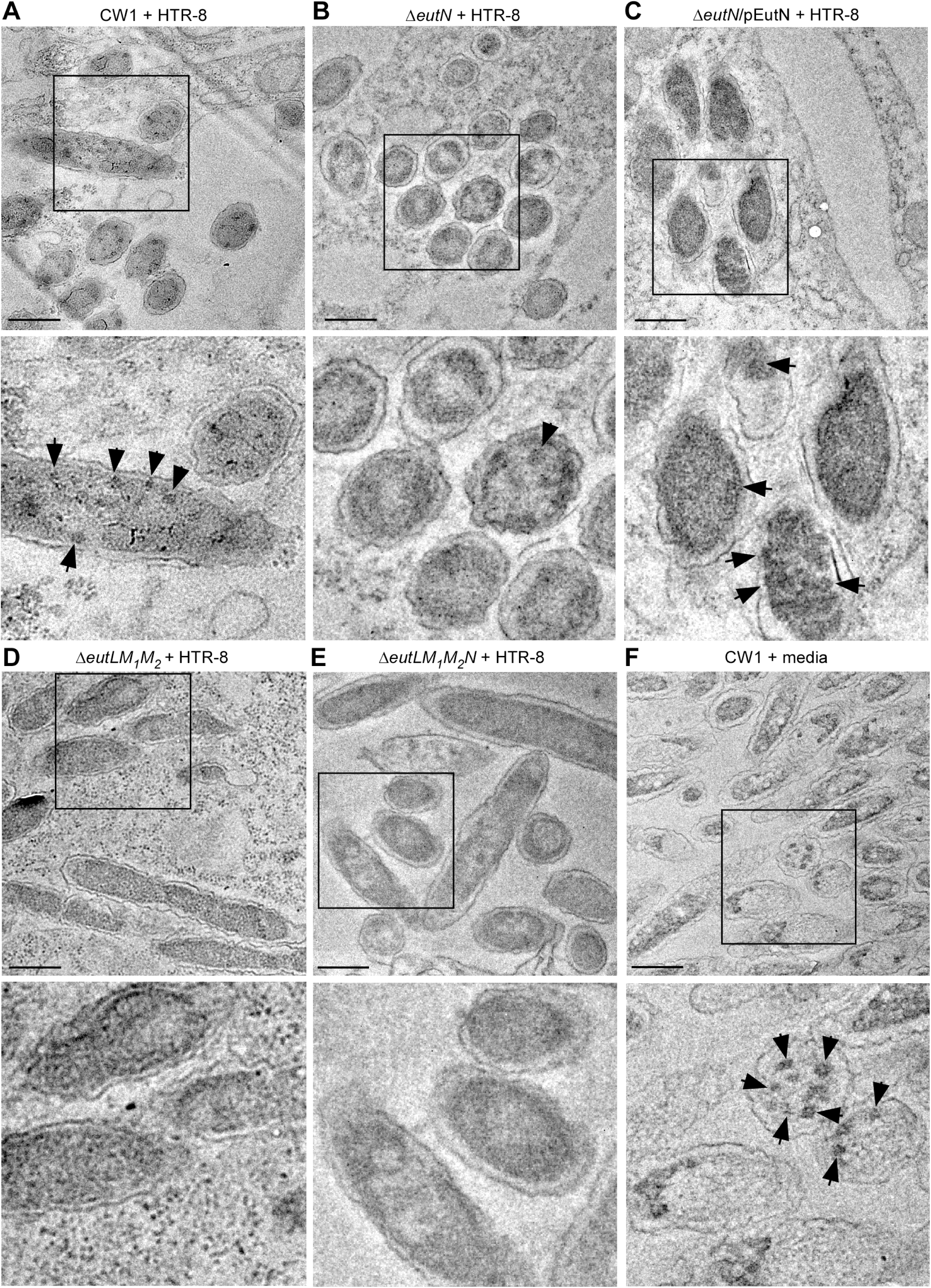
BMC formation in placental cells. **(A-F)** HTR-8/SVneo cells seeded at 3 × 10^6^ cells/ml were infected with *F. nucleatum* CW1 at the multiplicity of infection (MOI) of 450 for 4.5 h then embedded into resin before sectioning; scale bars of 200 nm. Black arrows in all images mark BMC structures. Images are representative of observed phenotypes.

Altogether, the results demonstrate that EutL, EutM_1_, EutM_2_, and EutN of *F. nucleatum* serve as a molecular determinant of leading to the EA-induced assembly of BMCs that are critical for bacterial growth under conditions whereby fusobacteria must utilize EA as a carbon and nitrogen source.

### BMC formation is critical for *F. nucleatum* virulence

Since deletion of either *eutN*, *eutL*/*eutM_1_*/*eutM_2_*, or *eutL*/*eutM_1_*/*eutM_2_*/*eutN* severely affects BMC formation, we next proceeded to determine whether the Δ*eutN* mutant has a virulence phenotype in a mouse model of preterm birth, using an experimental setup we tested out previously for examining other potential virulence factors (26, 29). We utilized groups of five pregnant CF-1 mice that were infected via tail vein injection on day 16/17 of gestation with ∼5.0 ×10^7^ CFU of the parent or Δ*eutN* mutant strain and monitored pup recovery and pup survival over time (Fig. 6A). Remarkably, the Δ*eutN* mutant showed a significant attenuation in virulence, with roughly 60% of pups being born alive by the end of experimentation, in contrast to the parent strain (CW1) which caused only 20% of pup survival (P < 0.001). Of note, no significant defects in growth and propagation of fusobacteria were observed in the Δ*eutN* mutant, as compared to the parent strain when fusobacterial cells were grown in rich media (Fig. 6C), a condition that was used to prepare bacterial cultures for infection. We conclude that BMC formation is critical for fusobacterial virulence in the mouse host.

**Figure 6:**
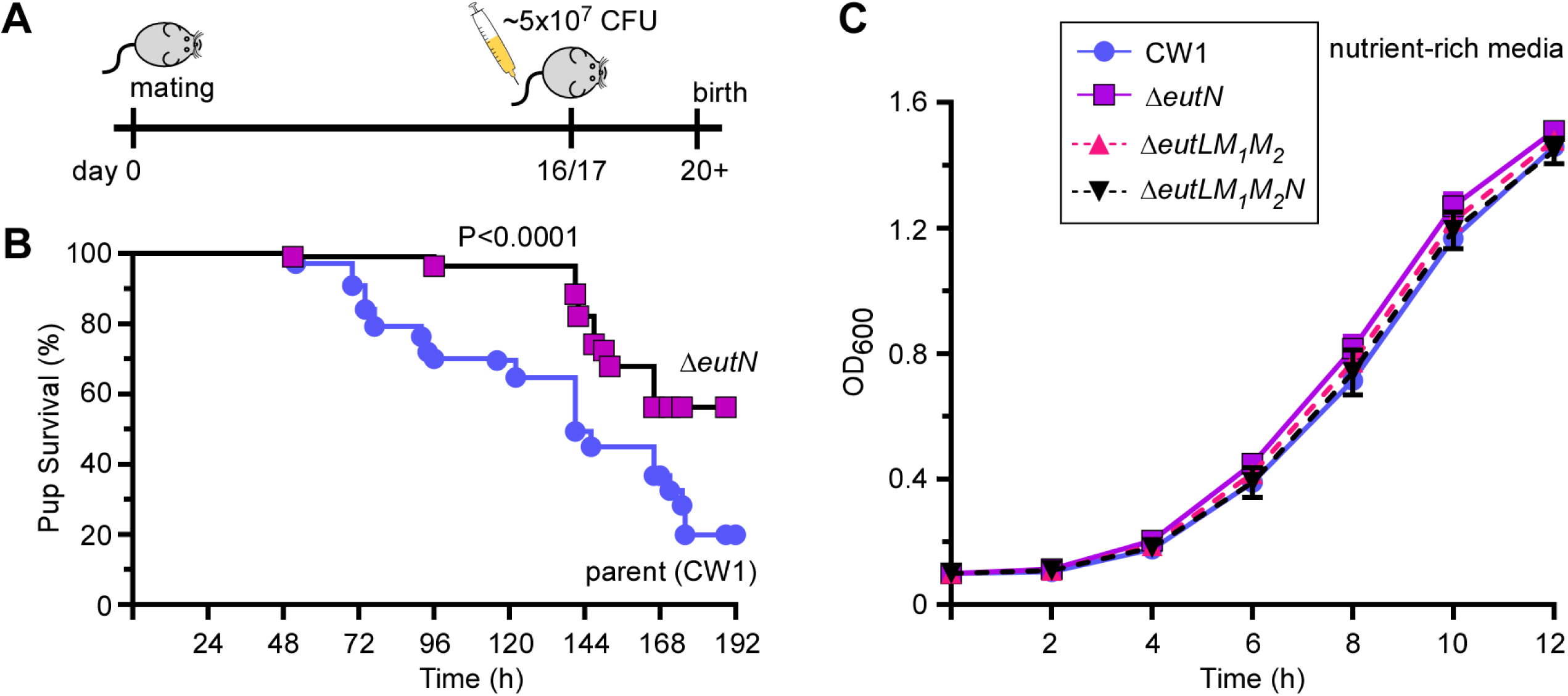
Virulence attenuation of the *eutN* mutant. **(A-B)** A group of 5 pregnant mice was infected with ∼5 × 10^7^ CFU of the parent or Δ*eutN* strain via the tail vein on day 16 or 17 of gestation, and pup survival was recorded. This experimented was repeated independently 3x. The statistical differences were analyzed by Mantel-Cox (*** P<0.001). **(C)** Cell growth of indicated strains in rich media were monitored by OD_600_.

## DISCUSSION

*F. nucleatum* is one of the key inhabitants of the human oral cavity that can migrate to and colonize multiple distal organs, promoting the development of several extra-oral diseases, most notably colorectal cancer and preterm birth (20). Our previous studies of fusobacterial pathogenesis employing targeted gene deletion mutants suggest that the metabolic versatility of *F. nucleatum* may contribute to its adaptability to different environments within its animal hosts (29). This is the trait that must be essential for a commensal organism to cause diseases in multiple organs that present unique challenges in varied cellular environments. Here, we present an important advance in support of this hypothesis by demonstrating that fusobacteria form specific metabolic organelles, BMCs, in response to environmental ethanolamine and that these BMCs serve as a major determinant of this pathobiont to maintain its virulent potential within its host.

Like many gastrointestinal bacteria, *F. nucleatum* harbors a large *eut* gene cluster, coding for 17 putative EA utilization (Eut) proteins and two hypothetical proteins, with the adjoining *folK* and *folP* genes seemingly a part of the *eut* transcriptional units (Fig 1A). We demonstrated that when EA is provided as a major carbon and nitrogen source, expression of many *eut* genes is significantly elevated, concomitant with increased BMC formation (Fig. 1). Interestingly, expression of the *eut* gene cluster appears to be bimodal, with *eutS*, *eutP*, *eutV*, and *eutW* genes down-regulated in the EA-conditioned media and the remaining genes starting from *eutA* highly expressed, including the genes coding for BMC shell proteins (*eutLM_1_M_2_N*), (Fig. 1D). This bimodal pattern of expression of *eut* genes is also mirrored in the gene expression analysis of the *eutV* mutant (Fig. 2A). The differential expression of *eutS* and *eutLM_1_M_2_N* suggests that EutS might not be part of the BMC structures in *F. nucleatum* under the tested experimental conditions. Consistent with this, mass spectrometry did not identify EutS among proteins associated with isolated BMCs (Table S2). Nonetheless, we cannot rule out a role of EutS in the process of BMC formation since we were unable thus far to obtain an *eutS* mutant via our conventional gene deletion method.

A key question our study addressed is the identity of the molecular components of *F. nucleatum* BMCs. Our biochemical and genetic analyses so far suggest that EutL, EutM_1_, EutM_2_, and EutN make up the shell of *F. nucleatum* BMCs: they are present as components enriched in the isolated BMCs, and the genetic disruption of EutLM_1_M_2_N severely affects BMC formation (Figures 3-5; Table S2). It is intriguing that *F. nucleatum* encodes two copies of EutM (EutM_1_ and EutM_2_), which are strikingly similar to *E. coli* EutM (30), except that EutM_1_ has an extra α-helix at the C-terminus as predicted by AlphaFold (31, 32) (Fig. S1). Moreover, while no EutK homologs are found in *F. nucleatum*, its *eut* gene cluster encodes two hypothetical proteins (RS2810 and RS2820). Whether these hypothetical proteins participate in BMC formation and if so, in what capacity remains to be investigated.

Significantly, we also demonstrated that BMCs are formed when fusobacteria are exposed to human placental cells in culture, an experimental condition that leads to the attachment and invasion of placental cells by fusobacteria (Fig. 5). In this context, it is important to note that fusobacterial BMCs are also observed when *F. nucleatum* was grown in Roswell Park Memorial Institute (RPMI) media supplemented with 5% fetal bovine serum (FBS) – the medium used for culturing the placental cells (Fig. 5F). The efficient assembly of fusobacterial BMCs observed in the RPMI/FBS culture medium alone might be mediated by EA and phosphatidyl-EA present in FBS (33) or some other yet unknown component that acts as an inducer of the eut genes and BMCs. Whatever the mechanism might be, we infer that fusobacteria are capable of assembling BMCs under conditions that mimic an environment pertinent to fusobacterial encounter of one of its target host tissue, the placenta, where it induces adverse pregnancy outcomes.

The induction of BMC formation and *eut* gene expression in response to EA suggested that these processes are transcriptionally regulated. Indeed, we showed that the response regulator EutV of the EutV-EutW pair constituting an EA-responsive TCS that is encoded within the *eut* locus modulates not only transcription of the *eut* locus in response to EA (Fig. 7A) but also the transcription of many additional genes outside of the *eut* gene cluster under a nutrient-rich condition (Fig. 2 and Fig. 7B). *F. nucleatum* EutV is annotated as a Fis family transcriptional regulator (biocyc.org), with Fis (factor for inversion <u>s</u>timulation) previously shown to be a regulator of metabolism in *E. coli* (34). Interestingly, as predicted by AlphaFold, EutV harbors an ANTAR domain (ANTAR for <u>A</u>miR and <u>N</u>asR transcription <u>a</u>ntitermination regulators), which is an RNA-binding domain identified in many response regulators of the two-component systems (35). Notably, as a representative member of the ANTAR-containing response regulators, the *E. faecalis* EutV regulates expression of *eut* genes (36), via its interaction with terminator elements located in the untranslated regions preceding *eutP*, *eutG*, *eutS*, and *eutA* (37). Whether or not a similar regulatory mechanism is operative in the case of the ANTAR domain in *F. nucleatum* EutV remains to be examined. It is worth mentioning that expression of *eut* genes appears to be cross-regulated by some other two-component systems and transcriptional regulators at certain conditions and cell growth. When cells are grown to mid-log phase in a nutrient-rich medium, deletion of either *carR*, coding for the response regulator CarR of the TCS CarRS or *modR*, encoding the response regulator ModR of the TCS ModRS, enhances expression of many *eut* genes (24, 25). In the early stationary phase, in mutants devoid of either the RNA-binding protein KhpA or KhpB, expression of 13 *eut* genes is downregulated (38).

**Figure 7:**
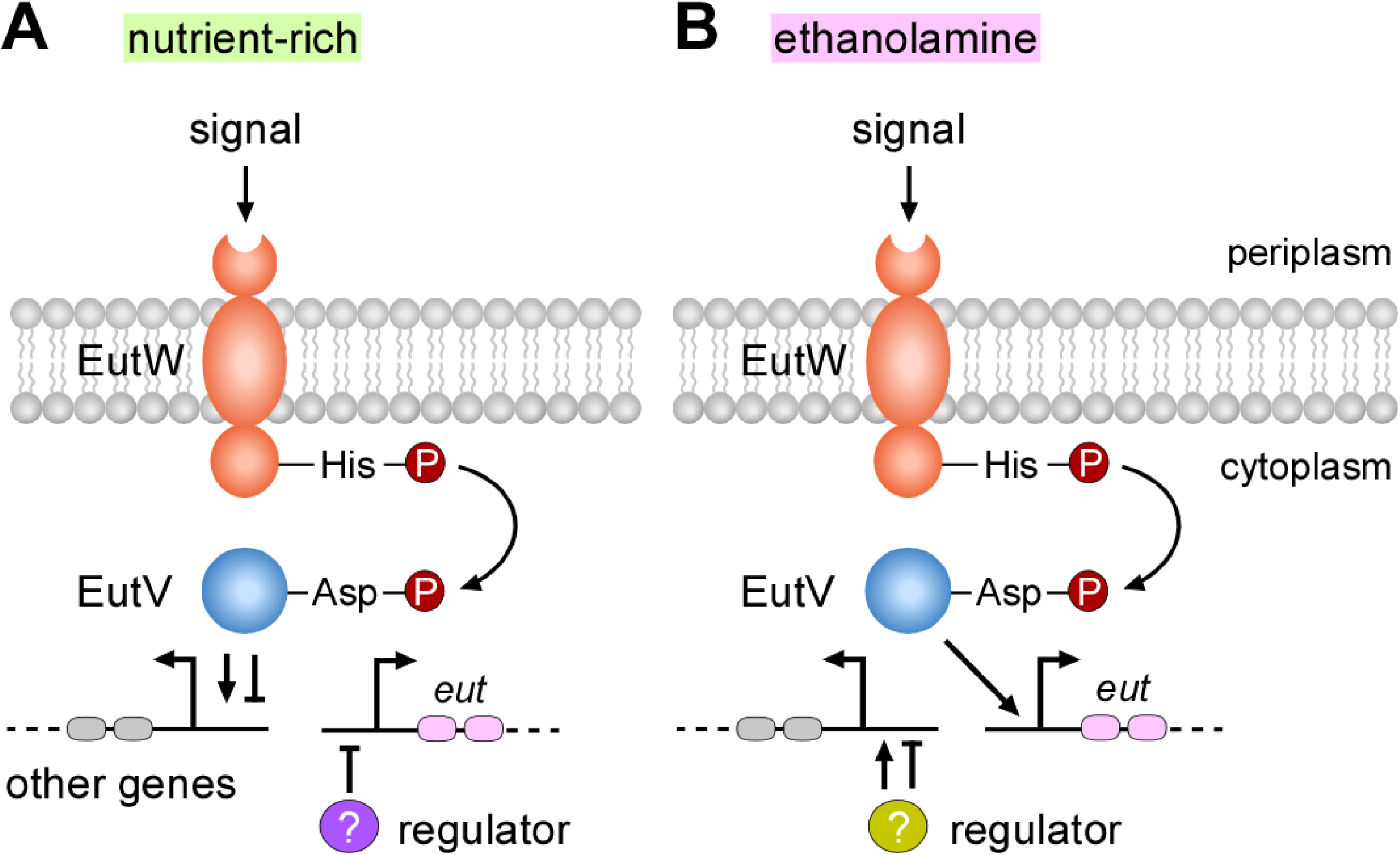
A working model of gene expression regulated by the two-component transduction system EutVW in *F. nucleatum*. **(A)** In nutrient-rich conditions, a specific signal(s) is recognized by the sensor kinase EutW, leading to autophosphorylation at a conserved histidine residue. The phosphate is transferred to a conserved aspartate residue of the response regulator EutV. Phosphorylation of EutV triggers the transcriptional regulation response of EutV, resulting in the activation or repression of various genes. It is unclear if EutV also regulates expression of a regulator that represses expression of *eut* genes in these conditions. **(B)** In the presence of environmental ethanolamine, which may act as a signal that activates EutW, the response regulator EutV becomes activated, leading to the upregulation of *eut* genes. In this condition, it is unclear if EutV and/or other regulators controls expression of other genes.

Perhaps more importantly, we demonstrated here that the ability of *F. nucleatum* to form BMCs is critical to its virulence potential in its host by showing that the *eutN* mutant that is defective in BMC formation is attenuated in the mouse model of fusobacterial infection causing preterm birth (Fig. 6). In this context, it is prudent to recall the striking clinical observation that the level of EA increases in the homogenates of human placental villi sampled during the third trimester, in parallel with a corresponding decrease in phosphor-ethanolamine, a precursor of EA (39). These observations lead us to posit that the ability of *F. nucleatum* to utilize environmental EA provides fusobacteria a physiologically significant advantage to adapt to specific niches such as the placenta and the colon, in spite of the changing and challenging environments that fusobacteria encounter along their long journey from the oral cavity to those distal organs.

## MATERIALS AND METHODS

### Bacterial Strains, Plasmids, and Media

Bacterial strains and plasmids used in this study are listed in *Supplemental Material* (*SM*) Table S3. *F. nucleatum* cells were grown in tryptic soy broth supplemented with 1% Bacto Peptone plus 0.05% cysteine (TSPC) or TSPC agar plates under anaerobic conditions (10% CO_2_, 10% H_2_, and 80% N_2_). *Escherichia coli* strains were grown in Luria broth (LB). When necessary, chloramphenicol and thiamphenicol were added to culture media at a concentration of 15 µg/mL and 5 μg/ml (Fisher Scientific), respectively. All reagents were purchased from Sigma-Aldrich and unless noted otherwise.

Genetic manipulation in *F. nucleatum*.

(i) *Generation of non-polar, in*-*frame deletion mutants* – Generation of single (Δ*eutV* or Δ*eutN*) and triple (Δ*eutLM_1_M_2_*) deletion mutants was performed according to a published protocol (26, 29). Briefly, 1-kb flanking regions upstream and downstream of *eutV*, *eutN* or *eutLM_1_M_2_* were PCR-amplified with appropriate primers (Table S4) and cloned into the deletion vector pCM-GalK (27), generating the corresponding deletion vectors pΔ*eutV*, pΔ*eutN*, and pΔ*eutLM_1_M_2_* (Table S3). The generated plasmids were electroporated into *F. nucleatum* CW1 cells, plasmid integration into the bacterial chromosome and producing integrant strains that were selected by thiamphenicol-containing TSPC agar plates. The integrant strains were grown in media without antibiotics to allow the second homologous recombination event. This led to plasmid excision that formed wild-type or mutant alleles, which were then selected on agar plates containing 2-deoxy-D-galactose and verified by PCR and DNA sequencing.

To create a quadruple mutant lacking *eutLM_1_M_2_N* (Δ*eutLM_1_M_2_N*), the Δ*eutLM_1_M_2_* deletion mutant and the deletion vector pΔ*eutN* (Table S3) were used in the deletion procedure mentioned above.

(ii) *Generation of complementing vectors* – To generate pYWC1, the 321-bp promoter region of the ribosomal RNA small subunit methyltransferase A (*rsmA*) in *F. nucleatum* ATCC23726 was PCR-amplified using appropriate primers (Table S4). The amplicon was treated with SacI and NdeI restriction enzymes prior to be ligated overnight at 16°C into pCWU6 precut with the same enzymes. The generated plasmid was introduced into *E. coli* DH5α for preparation of plasmid DNA, which was used for confirmation by DNA sequencing.

To generate pEutN, primers com-eutN-F and com-eutN-R were used to PCR-amplify the *eutN* coding sequence, with chromosomal DNA of *F. nucleatum* ATCC 23726 as a template, while appending BamHI/KpnI sites for cloning into pYWC1 precut with restriction enzymes BamHI/KpnI. The resulting vector, confirmed by DNA sequencing, was electroporated into the Δ*eutN* mutant strain.

### Bacterial growth assay

Overnight cultures of various *F. nucleatum* strains grown in TSPC were used to inoculate fresh cultures in minimal media with a starting optical density at 600 nm (OD_600_) of 0.1. Cell growth in an anaerobic chamber was monitored by OD_600_. The minimal media was made of 5x M9 salts (64 g of Na_2_HPO_4_•7H_2_O, 15 g of KH_2_PO_4_, 2.5 g of NaCl, and 5 g of NH_4_Cl in 1L (40), 1% Bacto peptone, 0.5 M MgSO_4_, 0.1 mM CaCl_2_, and 0.05% cysteine. Various concentrations of ethanolamine (Fisher Scientific) were added when necessary.

### RNA extraction

Total RNA of various *F. nucleatum* strains was extracted utilizing RNeasy Mini Kits (Qiagen) in accordance with the manufacturer’s protocol, with some modifications. Briefly, cell pellets harvested from 3-ml bacterial cultures were suspend in 200 µl of RNA-free 10mM Tris-EDTA, and the suspensions were transferred to a chilled Fast Protein Tube (Qbiogene) containing 700 µl of RNeasy lysis buffer (RNeasy Mini Kit, Qiagen) and 7 µL of β-mercaptoethanol (Thermo Fisher Scientific). Cells were lysed via Ribolyser (Hybaid), and supernatants were collected via centrifugation at 13,000 x g for 5 min at 4°C. RNA was then purified from the supernatants accordingly, and purified RNA was treated with deoxyribonuclease (DNase; Qiagen) and cleaned by using a RNeasy clean up kit (Qiagen).

### Gene expression analysis

For qRT-PCR, ∼100 ng of extracted RNA was used for complementary DNA (cDNA) synthesis using iScript Supermix (Bio-Rad) according to the manufacturer’s protocol. qRT-PCR was performed using a CFX96 Real-Time System (Bio-Rad), with reactions prepared using a SYBR Green PCR Master Mix (Bio-Rad) with appropriate primers (Table S4). Changes in gene expression were calculated as previously described (25).

For RNA-seq analysis, cDNA libraries were prepared, and sequencing was performed as previously reported (25). Genes with a log2 (fold change) above +1.0 or below −1.0 were considered to be differentially transcribed under the examined conditions.

### BMC isolation

Overnight cultures of various *F. nucleatum* strains in TSPC were used to inoculate fresh cultures (1:10 dilution), which were grown to mid-log phase. 8-ml aliquots of these cultures were transferred to 400 ml of minimal media supplemented with 50 mM of ethanolamine or without. The resulting cultures were grown overnight (14-16 h), and BMCs were isolated according to published protocols (8, 41). Briefly, harvested cells were washed in 20 ml of Buffer A (50 mM Tris, 500 mM KCl,12.5 mM MgSO_4_), followed by centrifugation at 5,000 x g for 15 min. Washed cells were resuspended in 10 ml of 60% bacterial protein extraction reagent (BPER) in Buffer A supplemented with 1 mM PMSF, 1 mg/ml lysozyme, and 10 µg/ml DNase and rocked at 30°C for 1 h. Supernatants were separated from cells by centrifugation at 12,000 x g for 8 min, and the supernatants were spun at 20,000 x g for 40 min to harvest BMCs. Crude BMCs were resuspended in 50 µl of chilled Buffer B (50mM Tris, 50mM KCl, 5mM MgCl_2_). BMC suspensions were centrifuged at 12,000 x g to remove contaminants.

### Mass spectrometry analysis of BMCs

Obtained BMC samples in lysis buffer (100 µl, 12 mM sodium lauroyl sarcosine, 0.5% sodium deoxycholate, 50 mM triethylammonium bicarbonate (TEAB)) were heated for 5 min. The samples were treated with tris(2-carboxyethyl) phosphine (10 µl, 55 mM in 50 mM TEAB, 30 min, 37°C), followed by treatment with chloroacetamide (10 µl, 120 mM in 50 mM TEAB, 30 min., 25°C in the dark). They were then diluted 5-fold with aqueous 50 mM TEAB and incubated overnight with Sequencing Grade Modified Trypsin (1 µg in 10 µl of 50 mM TEAB; Promega, Madison, WI) following which an equal volume of ethyl acetate/trifluoroacetic acid (TFA, 100/1, v/v) was added. After vigorous mixing (5 min) and centrifugation (13,000 x g; 5 min), the supernatants were discarded, and the lower phases were dried in a centrifugal vacuum concentrator. The samples were then dissolved in acetonitrile/water/TFA (solvent A, 100 μL, 2/98/0.1, v/v/v) and loaded onto a small portion of a C18-silica disk (3M, Maplewood, MN) placed in a 200 µl pipette tip. Prior to sample loading, the C18 disk was prepared by sequential treatment with methanol (20 µl), acetonitrile/water/TFA (solvent B, 20 µl, 80/20/0.1, v/v/v) and finally with solvent A (20 µl). After loading the sample, the disc was washed with solvent A (20 µl, eluent discarded) and eluted with solvent B (40 µl) (42). The collected eluent was dried in a centrifugal vacuum concentrator.

The eluants were reconstituted in water/acetonitrile/FA (solvent E, 10 µl, 98/2/0.1, v/v/v), and aliquots (5 µl) were injected onto a reverse phase nanobore HPLC column (AcuTech Scientific, C18, 1.8 μm particle size, 360 μm x 20 cm, 150 μm ID), equilibrated in solvent E and eluted (500 µl/min.) with an increasing concentration of solvent F (acetonitrile/water/FA, 98/2/0.1, v/v/v: min./% F; 0/0, 5/3, 18/7, 74/12, 144/24, 153/27, 162/40, 164/80, 174/80, 176/0, 180/0) using an Eksigent NanoLC-2D system (Sciex (Framingham, MA)). The effluent from the column was directed to a nanospray ionization source connected to a high-resolution orbitrap mass spectrometer (Q Exactive Plus, Thermo Fisher Scientific) acquiring mass spectra in a data-dependent mode alternating between a full scan (*m/z* 350-1700, automated gain control (AGC) target 3 × 10^6^, 50 ms maximum injection time, FWHM resolution 70,000 at m/z 200) and up to 10 MS/MS scans (quadrupole isolation of charge states ≥ 2, isolation width 1.2 Th) with previously optimized fragmentation conditions (normalized collision energy of 32, dynamic exclusion of 30 s, AGC target 1 × 10^5^, 100 ms maximum injection time, FWHM resolution 35,000 at *m/z* 200). The raw data was analyzed in Proteome Discoverer 2.4 with appropriate chemical modifications.

### Placental cells co-culturing with *F. nucleatum*

Placental HTR-8/SVneo cells were grown in Roswell Park Memorial Institute (RPMI) supplemented with 5% fetal bovine serum (FBS) to ∼90% confluence in an incubator maintained at 5% CO_2_ and infected with individual *F. nucleatum* strains, grown to mid-log phase in TSPC, at a MOI of 450 for 4.5 h. Cells were gently washed with ice-cold phosphate-buffered saline (PBS) to remove unattached cells prior to being subjected to the embedding procedure described below.

### Electron microscopy

BMCs were analyzed by transmission electron microscopy (TEM), adapted from a published protocol (27). Briefly, a drop of isolated BMCs in suspension was placed onto carbon-coated TEM grids and wicked off with filtered papers after 10 min. BMCs on grids were stained twice with 2% uranyl acetate prior to electron microscopic analysis (FEI Tecnai 20 iCorr TEM).

For thin-section TEM of fusobacteria alone or fusobacteria cocultured with placental cells, samples were prepared according to a published protocol with some minor modifications (43). For cocultured samples, cells were fixed overnight in 4% paraformaldehyde in 0.2 M HEPES at 4°C and washed with 0.1 M HEPES for 15 min. Cells were then fixed with 2% OsO_4_ and 1.5% potassium ferricyanide in 0.1M phosphate buffer at 4°C and dehydrated in increasing concentrations of pure-grade ethanol (50-100%) for 15 min with centrifugation at 13,523 x g for 3 min followed by 100% acetone for 7 min. Fixed cells were embedded in 33% Spurr’s resin in acetone for 1 h, 66% Spurr’s Resin in acetone overnight, and 100% Spurr’s Resin for ∼6 h and polymerized for ∼72 h. For bacterial cells alone, sample preparation was performed similarly except that cells were fixed in 4% paraformaldehyde in 0.2 M HEPES at room temperature for 2 h and washed in 0.1 M phosphate buffer, followed by post-fixation in 1% OsO_4_ in 0.1 M phosphate buffer and washed again. All samples were thin sectioned using an UCT Ultramicrotome (Leica) and collected on Formvar-coated nickel grids. Bacterial samples were post-stained with 2% uranyl acetate, while cocultured samples were post-stained with 3% lead citrate prior to analysis using a T20 iCorr TEM.

For cryoEM, 3 µL aliquots of BMC samples were applied to glow-discharged Quantifoil R2/1 holey carbon grids (300 mesh, Ted Pella). After 3-s incubation, grids were plunge-frozen into liquid ethane using a Vitrobot Mark IV vitrification system (Thermo-Fisher Scientific). During freezing condition optimization, cryo-EM grids were screened on an FEI Technai F20 electron microscope operated at 200 kV. Optimal conditions were achieved with a 90-s glow-discharge using a PELCO easiGlow™ (air as gas source, target vacuum of 0.37 mbar, target current of 15 mA) and Vitrobot settings at 4°C, 100% humidity, 5.5 s blotting time, blotting force 2, and 0 s drain time. CryoEM grids were imaged on the same instrument as used during screening, with a TIETZ F415MP 16-megapixel CCD camera at a nominal magnification of 25,000 × (pixel size 4.3 Å/pixel).

### Mouse model of preterm birth

To compare the infectivity of Δ*eutN* with that of the parent strain, we employed a mouse model of preterm birth as previously reported (26, 29). Briefly, groups of six CF-1 (Charles River Laboratories) pregnant mice were infected with ∼5×10^7^ CFU of the parent or Δ*eutN* strain at day 16 or 17 of gestation via tail vein injection. Pup survival was recorded for the next 8 days. Statistical analysis was determined via the Mantel-Cox text, using GraphPad Prism.

## DATA AVAILABILITY

All data generated in this study are provided in this manuscript, including the Supplemental Material. The RNA-seq data were deposited in the NCBI Gene Expression Omnibus (GEO) database with the accession number of GSE280934.

## ACKNOWLEDGMENTS

We thank Drs. Chenggang Wu, Ju Huck Lee, and Luong T. Truc (UT McGovern Medical School) for providing reagents, Lucy Gao and Andrew Goring (UCLA) for technical assistance, and the Ton-That lab members for their critical input and discussion. Research reported in this publication was supported by the National Institute of Dental & Craniofacial Research of the National Institutes of Health under the Award Numbers DE032906 and DE026758 (to H.T.-T). Dana Franklin was supported by the Ruth L. Kirschstein National Research Service Award T32AI007323. Angela Agnew was supported by the UCLA Dentist-Scientist and Oral Health-Researcher Training Program, NIDCR grant T90DE030860. The content is solely the responsibility of the authors and does not necessarily represent the official views of the National Institutes of Health.

## AUTHOR CONTRIBUTIONS

D.S.F., Y.W.C, and H.T-T. designed research; D.S.F., Y.W.C, Y.C., M.W., T.T.L, A.A, A.L. performed experiments; and all authors analyzed data. D.S.F., A.D. and H.T.T. wrote the manuscript. All authors read and approved the final manuscript.

## DECLARATION OF INTERESTS

The authors declare no competing interests

